# A permissive role for dopamine in the production of vigorous movements

**DOI:** 10.1101/2022.11.03.514328

**Authors:** Haixin Liu, Riccardo Melani, Akhila Sankaramanchi, Ruoheng Zeng, Marta Maltese, Jenna R. Martin, Nicolas X. Tritsch

## Abstract

Dopamine is essential for the production of vigorous movements, but how dopamine modifies the gain of motor commands remains unclear. Here, we developed a dexterous motor task in which head-restrained mice self-initiate fast and large-amplitude lever pushes with their left forelimb to earn rewards. We show that this task is goal-directed and depends on cortico-striatal circuits in the hemisphere contralateral to the limb used to push the lever. We find that unilateral loss of midbrain dopamine neurons reduces the speed and amplitude of lever pushes, and that levodopa treatment rapidly restores motor vigor, consistent with parkinsonian bradykinesia. Photometry recordings of striatal dopamine levels indicate that the therapeutic efficacy of levodopa does not require phasic dopamine release. In dopamine-intact mice, optogenetic stimulation of midbrain dopamine neurons calibrated to mimic transients evoked by rewards is also insufficient to increase the speed and amplitude of forelimb movements. Together, our data show that phasic dopamine transients are unlikely to specify the vigor of forelimb movements online as they are being executed, and suggest instead that dopamine plays a permissive role in the selection and production of vigorous movements. Our findings have important implications for our understanding of how the basal ganglia contribute to motor control under physiological conditions and in Parkinson’s disease.

## Main Text

Ever since the discovery that the degeneration of midbrain dopamine (DA) neurons projecting to striatum underlies bradykinesia in Parkinson’s disease (PD), DA has become synonymous with motor vigor^1^. However, the mechanisms through which DA contributes to the speed and amplitude of voluntary movements are still debated^2-4^. Initial investigations suggested a permissive role for DA in allowing striatal circuits to function properly, since restoring DA signaling pharmacologically alleviates motor impairments in PD patients as well as in reserpinized animals, albeit incompletely^5-9^. More recently, DA neurons were shown to dynamically encode kinematic parameters and to phasically increase their activity at movement onset in proportion to vigor^10-14^, suggesting that rapid changes in striatal DA may dynamically control the gain of descending motor commands originating in cortex as they are routed through striatum to brainstem and spinal motor circuits for movement execution. However, studies employing optogenetics to test the ability of DA transients to invigorate movements on a moment-by-moment basis have yielded conflicting results^12-18^, possibly because of differences in the strength and duration of optogenetic manipulations, the sensitivity of experimental measures to movement speed and amplitude, and the extent to which movements commonly studied in rodents (e.g. licking, locomotion, lever pressing, etc.) depend on cortico-striatal circuits (and consequently on striatal DA levels) for their vigorous execution.

To elucidate how striatal DA contributes to vigor, we designed a dexterous motor task in mice inspired by tests used to measure bradykinesia in humans. PD patients typically show deficits in the production of fast, large-amplitude, self-initiated forearm and finger movements, which typically manifest unilaterally early in disease before progressing to other extremities and bilaterally^19, 20^. We therefore we trained head-restrained mice (N = 23) to grab a loaded lever with their left forepaw and to perform self-paced ballistic pushes following an uncued movement-free interval (**Fig. 1a–c;** see Methods). Inward pushes exceeding experimentally-set amplitude (mean: 4.1 ± 0.2 mm) and duration (< 0.5 s) criteria were rewarded with delayed (0.5 s) delivery of water from a lick spout. Well-trained mice learned to reliably produce rewarded pushes within a few seconds of uncued trial starts (**Fig. 1d–g**). Importantly, mice stopped pushing altogether if water was devalued (**Fig. 1h**), indicating that their behavior is goal-directed and therefore likely to engage cortico-striatal circuits^21^.

**Fig. 1:**
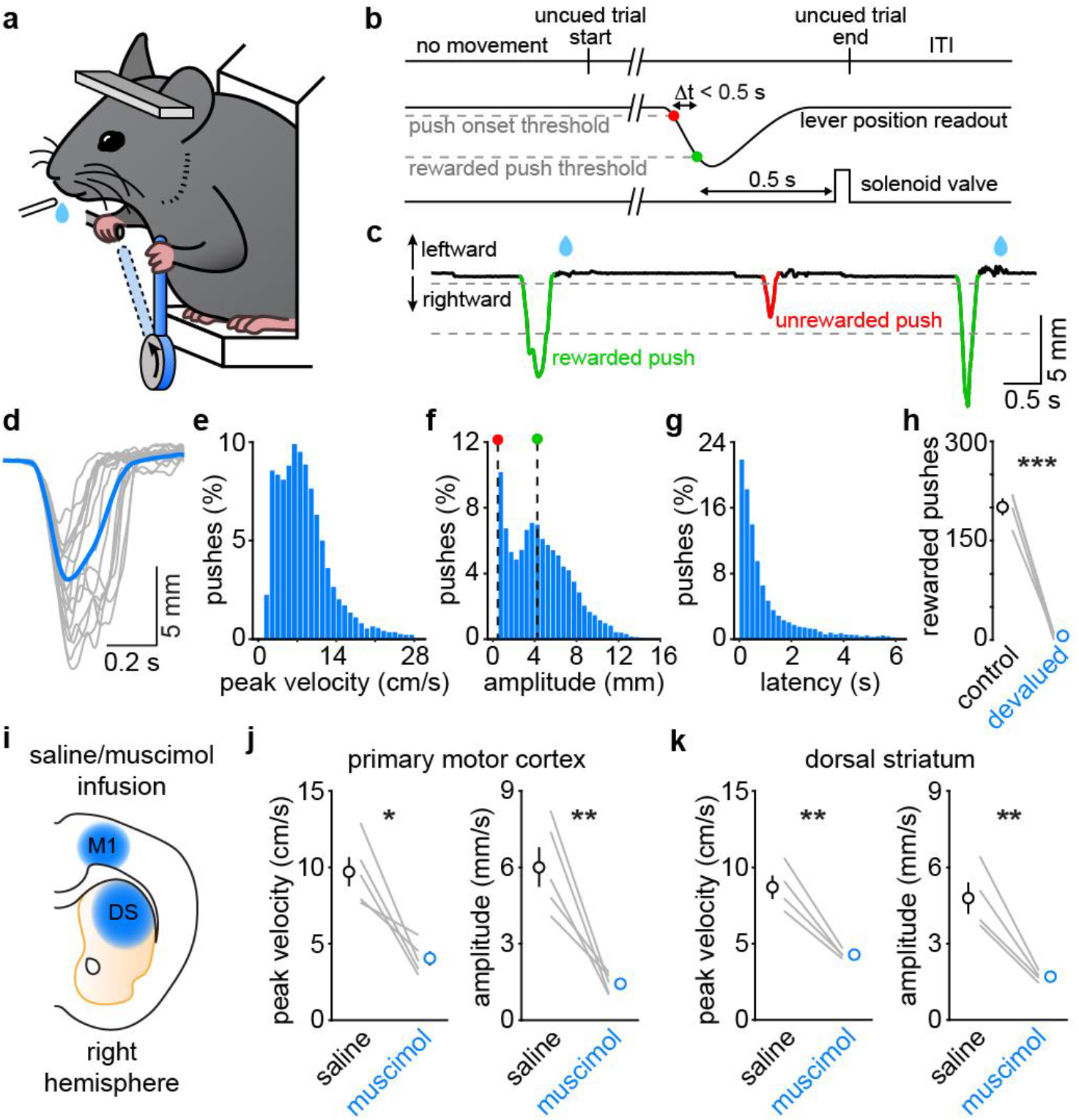
Self-paced lever task requires motor cortex and the basal ganglia. **a**, Experimental setup. **b**, Trial structure: mice must self-initiate a ballistic rightward lever push large enough to cross both thresholds (dashed gray lines) within 0.5 s to earn a delayed water reward. ITI: inter-trial interval. **c**, Example lever trace showing 3 self-paced rightward pushes. Reward schematized as blue drop. **d**, Example lever trajectory aligned to push onset from one mouse (gray: 15 randomly-selected pushes; blue: session mean). **e-g**, Probability distribution of peak velocity (**e**), maximum amplitude (**f**), and latency to reach maximum amplitude from uncued trial start (**g**) for all recorded lever pushes across 40 sessions in 18 mice. Dashed lines in **f** indicate push onset (red) and rewarded push (green) thresholds. **h**, Total number of reward collected in 4 mice at baseline and following reward devaluation. ***p=0.001, paired t-test. **i**, Schematic of mouse brain in coronal plane showing location of infusions in primary motor cortex (M1) or dorsal striatum (DS). **j**, Session medians of peak push velocity (*p=0.011; paired t-test) and amplitude (**p=0.007; paired t-test) following saline or muscimol infusion in M1 in 5 mice. **k**, same as **j** for DS (velocity p=0.006; amplitude p=0.010; paired t-test, N=4 mice). Lines in **h**, **j** and **k** reflect paired data for each mouse, and summary data are mean±SEM for each group.

To establish whether performance of this task indeed depends on cortico-striatal circuits, we acutely silenced neural activity in the forelimb region of primary motor cortex (M1; N = 5 mice) or dorsal striatum (DS; N = 4 mice) in the right hemisphere (i.e., contralateral to the limb used to push the lever) using local infusion of the GABA_A_ receptor agonist muscimol. In both cases, muscimol drastically impaired the mice’s ability to generate fast, large-amplitude lever pushes compared to saline infusions (**Fig. 1i–k**), confirming that M1 and DS participate in the production of vigorous forelimb movements in our task.

We next tested whether motor performance depends, like in humans, on the integrity of DA axons in the striatum contralateral to the limb under study. To do so, we compared mice (N = 8) before and after chronic unilateral lesion of ventral midbrain DA neurons (mDANs) in the right hemisphere with 6-hydroxydopamine (6-OHDA; **Fig. 2a**). Post-hoc histology confirmed near complete degeneration of DA axons in the right striatum (**Fig. 2b,c**). Lesioned mice continued to self-initiate lever pushes, although their movements were significantly slower and smaller in amplitude (**Fig. 2d–f**), consistent with parkinsonian bradykinesia. Importantly, restoring DA with a single low dose of the DA precursor levodopa effectively alleviated bradykinesia, temporarily returning the speed and amplitude of lever pushes to pre-lesion performance within minutes of administration and for the duration of the task (**Fig. 2d-f**). Together, these data confirm that DA is required for the production of vigorous contralateral limb movements, and indicate that performance in this task is dependent on contralateral DA signaling.

**Fig. 2:**
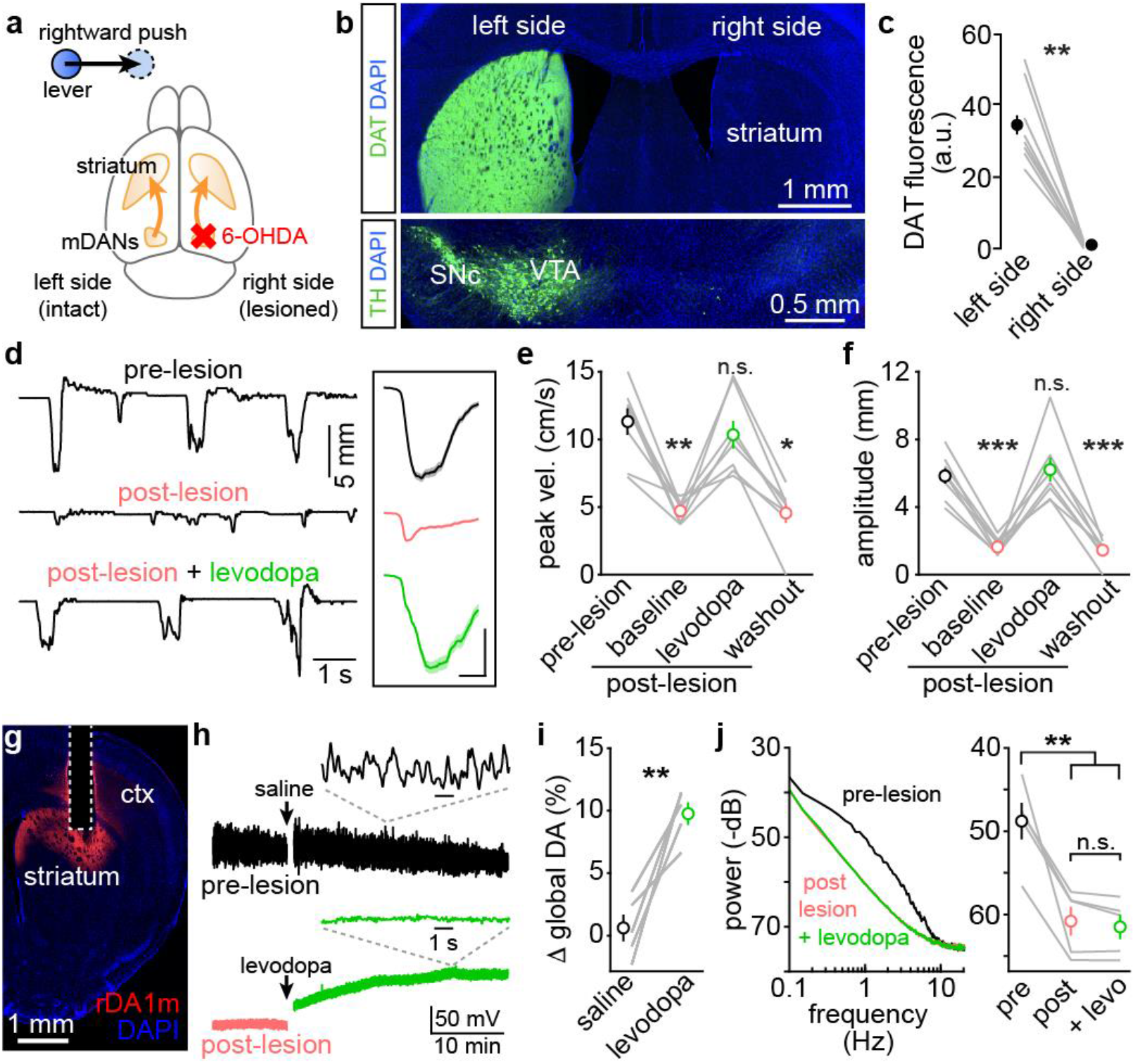
Striatal dopamine is required for the production of vigorous forelimb movements. **a**, Experimental setup; mDANs were lesioned contralaterally to the limb used to push the lever. **b**, Example fluorescence images of forebrain (*top*) and ventral midbrain (*bottom*) in the coronal plane stained for DAT and TH to label DA axons and cell bodies, respectively. SNc, substantia nigra pars compacta; VTA, ventral tegmental area. **c**, Quantification of DAT fluorescence in DS. **p=0.008, Wilcoxon, N=8 mice. **d**, Example lever traces before (*top*) and after (*middle*) mDAN lesion, and following levodopa treatment (*bottom*). Inset: push onset-aligned session averages (scale: 2 mm, 0.2 s). **e**, Median peak velocity of lever pushes across conditions. p<0.001, Friedman; Dunn–Šidák *post-hoc* comparisons between pre-lesion vs. post-lesion baseline (p=0.002), vs. levodopa (p=0.99), vs. washout (p=0.012). N=8 mice. **f**, Same as **e** for peak amplitude. p=0.002, Friedman; Dunn–Šidák: pre-lesion vs. post-lesion (p=0.022), vs. levodopa (p=1.000), vs. washout (p=0.006). **g**, Coronal section showing fluorescent DA sensor in DS. DAPI, nuclear stain; Ctx, cortex. **h**, Bleaching-corrected rDA1m signal in DS before (*top*) and after mDAN lesion (*bottom*) upon systemic administration of saline or levodopa. *Insets*: detailed view of intrinsic DA fluctuations. **i**, Changes in global rDA1m fluorescence evoked by saline and levodopa. **p=0.005, paired t-test, N=5 mice. **j**, *Left*, example power spectrum density of unprocessed rDA1m signal recorded from same mouse across conditions. *Right*, mean power in 0.5–4 Hz band. p<0.001, one-way ANOVA; Dunn–Šidák: pre-lesion vs. post-lesion (p=0.002), vs. post-lesion+levodopa (p=0.001), post-lesion vs. post-lesion+levodopa (p=0.99). Lines in **c**, **e**, **f**, **i** and **j** reflect paired data for each mouse, and summary data are mean±SEM for each group. ns: not significant.

The superior efficacy of levodopa over other therapies (including DA receptor agonists) in the symptomatic treatment of PD has been attributed to the ability of levodopa to restore phasic DA release from surviving DA axons or from serotonergic axons in striatum^22, 23^. To assess this, we imaged extracellular DA in DS using photometry of the fluorescent genetically-encoded DA sensor GRAB-rDA1m^24^ (**Fig. 2g**). Levodopa significantly elevated basal extracellular levels of DA in the DS of denervated mice (**Fig. 2h,i**). However, the brief periodic fluctuations in extracellular DA that were present in the striatum of the same mice prior to DA denervation were not restored (**Fig. 2h,j**), indicating that the therapeutic efficacy of levodopa does not require phasic DA release. Instead, our data suggest that some basal level of DA signaling is sufficient for cortico-striatal circuits to produce vigorous actions.

These data do not exclude the possibility that phasic elevations in DA also contribute to motor vigor by applying an additional ‘online’ gain function to motor commands, as they are relayed from cortex to brainstem and spinal circuits via the basal ganglia. To test whether phasically elevating DA can increase the vigor of forelimb movements, we optogenetically activated the cell body of channelrhodopsin-2 (ChR2)-expressing mDANs contralateral to the lever-pushing forelimb on a subset of trials (30%, interleaved design) with blue light (470 nm) during the first second of the uncued trial start (**Fig. 3a**), since well-trained mice initiate 52% of lever pushes during that time (**Fig. 1g**). We calibrated our stimulation parameters to produce increases in DA in DS as large as or slightly larger than those evoked by water rewards (**Fig. 3b,c**). Despite significantly elevating DA levels, however, ChR2 stimulation did not increase the speed or amplitude of lever pushes compared to trials without stimulation (baseline), or to sham stimulations, where red illumination (595 nm) was used instead of blue to control for the effects of light delivery to the midbrain (**Fig. 3d–f** and **Extended Data Fig. 1a,b**). To account for the fact that DA takes a few hundred milliseconds to modify the excitability of striatal projection neurons^25^, we limited our analysis to lever pushes initiated 1 s after stimulation onset but did not observe any difference either (**Extended Data Fig. 1c**). We also failed to observe notable effects on the speed or amplitude of lever pushes produced while DA levels are elevated (**Extended Data Fig. 1d**), or using stronger (10 mW) or weaker (1 mW) blue light intensities, excluding concerns that DA levels were too low or too high to reveal phasic effects on forelimb movement vigor (**Extended Data Fig. 1e,f**). In addition, unilateral optogenetic stimulation of mDANs in the same mice using identical parameters was sufficient to reinforce operant behavior (**Fig. 3g,h**), a hallmark of phasic DA^17^. Lastly, to confirm that our behavioral assay is sensitive to increases in the vigor of forelimb movements, we subjected the same mice to behavioral sessions in which the amplitude threshold needed to earn rewards suddenly increased mid-session. Consistent with goal-directed performance, mice significantly increased the speed and amplitude of lever pushes needed to obtain reward compared to sham sessions where the rewarded amplitude remained stable (**Fig. 3i–k**). Together, these results indicate that phasic elevation of DA is not sufficient to increase the vigor of ongoing forelimb movements.

**Fig. 3:**
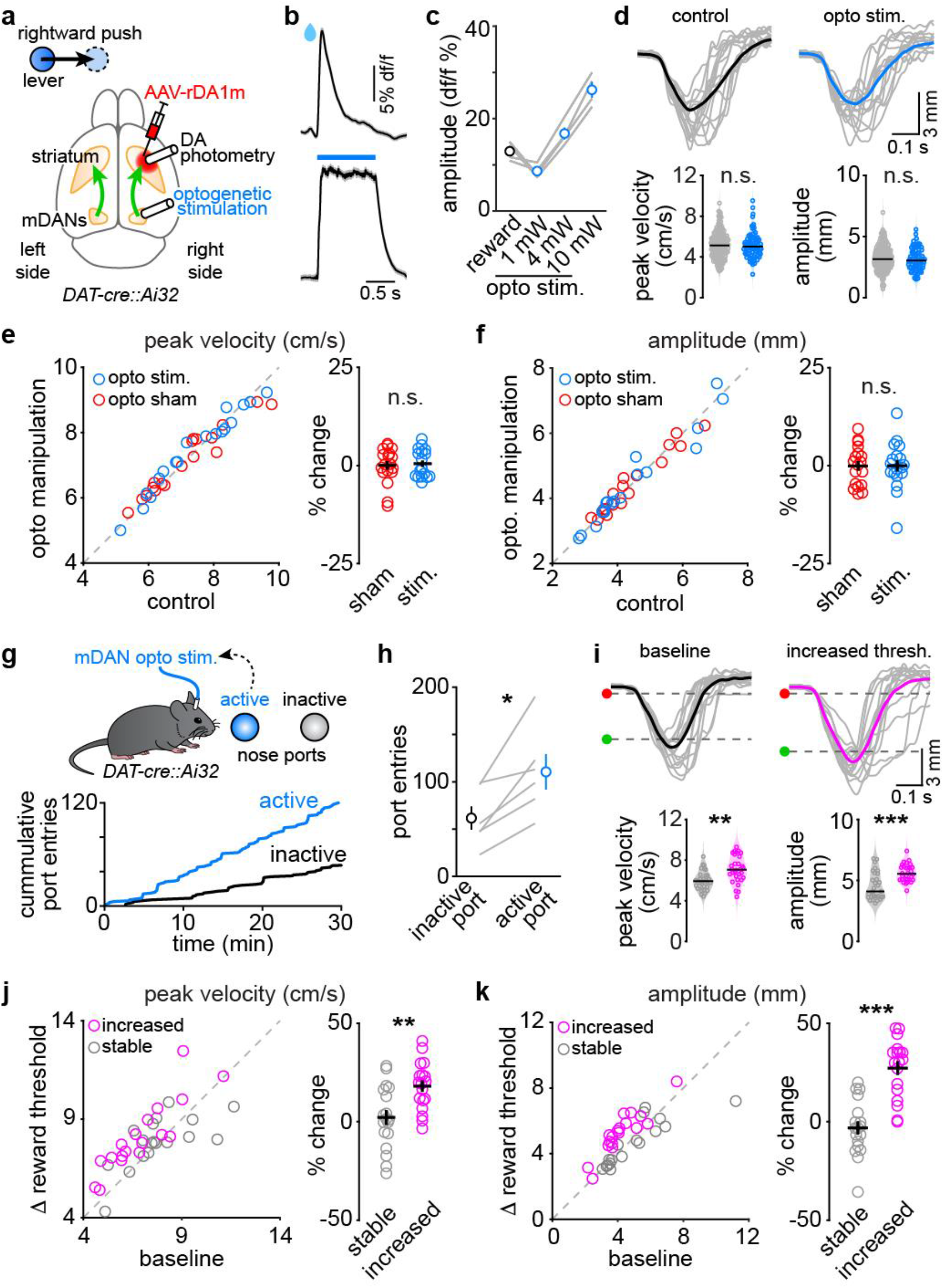
Phasic DA increase does not invigorate ongoing forelimb movements. **a**, Experimental setup: mDANs expressing ChR2 are activated with blue light (470 nm) in the ventral midbrain while DS DA levels are imaged with rDA1 m photometry. **b**, Example mean ±SEM rDA1m signal evoked by water delivery (*top*) and optogenetics mDAN activation (*bottom;* 4 mW, 10 ms pulses at 30Hz). **c**, Calibration of optogenetic DA elevation in DS relative to reward. **d**, *Top*, example lever push (color: session average; gray: 15 individual pushes) on trials with (*right*) and without (*left*) mDAN activation. *Bottom*, peak push velocities (p=0.82; Mann–Whitney) and amplitudes (p=0.58, Mann–Whitney) recorded during one session in one mouse. Black bar: session median. **e**, *Left*, scatter plot of median peak velocity in control trials (no stimulation) vs. trials with blue (mDAN stimulation) or red (sham stimulation) light. *Right*, % change in median peak velocity compared to control trials. p=0.72, Friedman. N=18 sessions in 6 mice. **f**, Same as **e** for amplitude. p=1.0. **g**, *Top*, experimental setup: unilateral mDAN stimulation triggered with nose poke in active port. *Bottom*, cumulative port entries for one mouse. **h**, Total number of port entries. p=0.01, paired t test. N=6 mice. **i**, Same as **d** before and after uncued increase in reward threshold. Velocity: p=0.003; amplitude, p<0.001, Mann–Whitney. **j,k**, Same as **e** and **f** comparing the maximum velocity (p=0.008) and amplitude (p<0.001) of rewarded lever pushes before and after reward threshold changes (magenta: increased; gray: unchanged). N=18 across the same 6 mice as in **d-f**. Summary data in **c**, **e**, **f**, **h**, **j** and **k** are mean±SEM for each group. ns: not significant.

In this study, we sought to clarify the role of phasic DA release in controlling the vigor of movements. To do so, we developed a behavioral assay that 1) depends on forebrain structures on which mDANs act, 2) precisely measures the speed and amplitude of movements, and 3) mimics PD’s characteristic levodopa-responsive distal bradykinesia that initially led to the idea that DA contributes to the production of fast movements. Consistent with clinical observations in PD patients, we found that levodopa acutely reinstates the speed and amplitude of contralateral forelimb movements in mice with unilateral loss of forebrain DA. Importantly, levodopa treatment restored global DA levels in striatum but not intrinsic DA transients, indicating that that sub-second fluctuations in DA are not necessary for the vigorous execution of volitional arm movements. In line with this, DA receptor agonists also alleviate bradykinesia in PD and restore vigorous movements in animals treated with the vesicular monoamine transporter antagonist reserpine, in which phasic release of all monoamines is abolished^5-8^.

In DA-intact mice, stimulation of mDANs using optogenetics failed to increase the speed or amplitude of voluntary forelimb movements, whether they were produced while DA levels were high, or in the seconds that followed. This negative finding did not stem from a failure to elevate striatal DA sufficiently as stimulation power was calibrated to match or exceed reward responses, nor was it due to a ceiling effect on vigor as mice readily increased the speed and amplitude of lever pushes to optimize reward collection. Our data therefore indicate that the speed and amplitude of ballistic arm movements are not directly determined by the levels of extracellular DA that exist in the basal ganglia when (or shortly before) motor commands are issued. As such, the reported phasic modulation of mDAN firing at movement onset may merely correlate with – but not specify – the vigor of ongoing actions. Indeed, many mDANs actually decrease their activity around the initiation of self-initiated movements such as locomotion, particularly when actions are not paired with reward^13^, ^17^, ^26^. Of the mDANs that increase their firing, few do so prior to movement onset, and the correlation between changes in activity and upcoming velocity and/or acceleration is weak^12^, ^13^, ^27^. In addition, the time-varying speed of reaching movements is encoded in the activity of M1 neurons and can be decoded to control the speed of prosthetic limbs in monkeys and humans^28-30^, suggesting that phasic release of DA is unlikely to play a central role in setting the gain of motor commands downstream of motor cortex while movements are being produced. In line with this, the neuromodulatory effects of DA are not immediate, as DA signals exclusively through G protein-coupled receptors^31^.

DA and the striatal neurons on which it acts have been extensively implicated in motivation – the incentivizing and energizing of motor actions in the pursuit of a goal^32, 33^. Motivation is the behavioral manifestation of several neural processes, ranging from action selection (e.g., bias for or against choosing an energetically-costly action given an expected outcome) to action initiation (e.g., changes in the threshold required to engage in a behavior) and action performance (e.g., gain modulation of motor commands). These processes can be difficult to dissociate using manipulations that produce supraphysiological elevations of DA over multiple behavioral epochs, and using session-averaged behavioral measures such as latency to reach a port, distance travelled, or number of licks or presses performed. Our optogenetic manipulations timed to the moment mice execute discrete forelimb movements indicate that DA is unlikely to invigorate behavior by controlling the moment-to-moment gain of descending motor commands (i.e. action performance). This is consistent with the observation that rewards do not progressively increase the speed and amplitude of lever pushes over the course of behavioral sessions, but failures to earn expected rewards (e.g., when the reward threshold is increased) do. Thus, the behaviorally motivating effects of DA may be primarily mediated at the level of action selection and/or action initiation. Indeed, compelling evidence supports a role for DA and striatal circuits in learning and deciding which action (including its vigor) to produce in a given context and when^4, 34, 35^. In light of this, parkinsonian bradykinesia may represent an implicit decision to move slowly (i.e. a selection problem) more so than an inability to energize task-appropriate motor commands issued by cortex (i.e. an execution problem)^36-41^.

## Methods

### Animals

All procedures were performed in accordance with protocols approved by the NYU Langone Health (NYULH) Institutional Animal Care and Use Committee (protocol #170123). Mice were housed in group before surgery and singly after surgery under a reversed light-dark 12-hour cycle (dark from 6 a.m. to 6 p.m.). Food and water were provided *ad libitum*, except when water-restricted to incite consumption of water rewards. Experiments were carried out using both male and female mice maintained on a C57BL/6J background (Jackson Laboratory; #000664) at 8–52 weeks of age. For optogenetic experiments, we genetically expressed ChR2 in mDANs by crossing *Dat-ires-Cre* (Jackson Laboratory; #006660) and *Ai32* (Jackson Laboratory; #012569) mouse lines to obtain offspring heterozygous for both transgenes.

### Surgeries

Mice were anesthetized with isoflurane and placed in a stereotaxic apparatus on a heating pad. Ketoprofen (10 mg/kg in saline) was injected subcutaneously. After exposing and cleaning the skull under aseptic conditions, coordinates (given in mm from bregma) were marked on the skull in the right hemisphere (M1: AP +0.3, ML +1.5; DS: AP 0, ML +2.8; SNc: AP −3.14, ML +1.42) and were either used to guide craniotomies or covered with Kwik-Cast (WPI) before applying dental cement (C&B metabond, Parkell) for future injections. A custom titanium headpost was attached to the skull in the horizontal plane posterior to lambda using a 3D printed, negative pressure-based holder and dental cement. To allow for drug infusions in DS, a cannula (P1 Technologies; C315GS-5) was implanted at a depth of 2.05 mm from dura. To image striatal DA by photometry, 200 nl of an adeno-associated virus (AAV) encoding the red-shifted GRAB-DA sensor rDA1m (AAV9-hSyn-rDA1m; titer: 6.37 e+12; Vigene Biosciences) was infused in DS at a depth of 2.05 mm below dura using a micro-syringe pump (KD Scientific; Legato 111; rate: 100 nl/min) fitted with a Hamilton syringe (1701 N) connected to a pulled glass injection micropipette (Drummond Wiretroll II; 100 μm tip) via PE tubing filled with mineral oil. A 4 mm-long fiber optic cannula (400 μm core, 0.5 NA, 1.25 mm ferrule, >80% light transmission efficiency; RWD Life Science Inc.) was subsequently implanted 0.2 mm above the injection site. To optogenetically stimulate mDANs, a 6 mm-long fiber optic cannula was implanted at a depth of 4.0 mm from dura in *Dat-ires-Cre::Ai32* mice. Implants were cemented to the skull using C&B metabond. Mice were allowed to fully recover in their home cage for at minimum 1 week before initiating behavioral training.

### Self-paced lever-pushing task

Behavioral rigs consist of a soundproof chamber, an aluminum headpost holder, a 3D-printed enclosure with foam cushion to hold the mouse, a 3D-printed lever with handle attached to an analog rotary encoder (US Digital; MA3 magnetic shaft encoder) loaded with a pair of magnets (one attached to the lever and another circular magnet affixed to the floor of the chamber) and a water delivery tube connected to a quiet solenoid valve (Lee Company; LHQA0531220H). Task parameters were controlled by an Arduino Due micro-processor running an open-source package (Bpod, v0.5, Sanworks). The voltage signal provided by the lever’s rotary encoder was handled by another Arduino Due; its signal was smoothed online (10 ms median filter) and threshold crossings were determined online using custom Arduino programs and a MATLAB graphical-user interphase. Analog and digital signals (e.g., lever position, synchronization signals, TTLs) were digitized at 10 kHz using a National Instruments data acquisition board (PCIe-6353) and breakout terminal block (BNC-2090A) and recorded using Wavesurfer (Janelia; v1.0.6). Each trial started after a 0.3 s period during which the lever voltage remained below a threshold set near the resting position to detect ‘push onsets’. Trials ended after mice completed a successful push and obtained a water reward or 30 s elapsed, whichever occurred first. A successful lever push was defined as any rightward lever push that crossed ‘push onset’ and ‘rewarded push’ thresholds (mean amplitude across mice: 4.1 ± 0.2 mm) within 0.5 s. The ‘rewarded push’ threshold was set to be large enough for rewarded pushes to be deliberate and effortful, but well-within the range of amplitudes that mice can produce (see Fig. 1f). Water rewards (~ 5 μl) were delivered 0.5 s after crossing of the ‘rewarded push’ threshold to separate forelimb from consummatory movements. The end of each trial was followed by a 1 s inter-trial interval to allow for water consumption. To promote learning and incite behavioral performance, mice had their access to water restricted to the recording rig. Their weight was monitored daily to ensure it remained above 85% of their original body weight. Mice were introduced to the task gradually; they were first habituated to head-fixation and water collection from the lick spout, before being trained to push the lever for reward with increasingly larger pushes. Mice reached expert performance (~200 rewarded trials in 30 min or less) after approximately 14 daily training sessions. To probe whether performance is goal-directed, we devalued water rewards by providing mice with free access to water 12 h prior to testing. To probe whether expert mice can adjust the speed and amplitude of lever pushes, we suddenly increased the ‘rewarded push’ threshold by ~10% on some sessions on trial 80.

### Muscimol inactivation

To acutely silence neural activity in M1 or DS, we slowly infused 200 nl of the GABA_A_ receptor agonist muscimol (Tocris; 2 μM in saline) into the target area using a micro-syringe pump (KD Scientific, Legato 111; 50 nl/min) while mice were briefly and lightly anesthetized with isoflurane in a stereotaxic frame. For DS inactivation, we used an infusion cannula (P1 Technologies; C315IS-5) through a chronically-implanted guide cannula (see above). For M1 inactivation, we made two 100 nl injections at 400 μm and 900 μm below dura using a sharp glass pipette (<50 μm tip) through a small craniotomy made over M1 the day prior. Infusion cannulas and glass needles were left in place for 10 min post-infusion. Mice were then returned to their home cage, where they emerged from anesthesia within minutes. Behavioral sessions started 10 min later. Control sessions using saline instead of muscimol were performed using the same procedure.

### Unilateral mDAN lesions

Mice were given free access to water for at minimum 2 days ahead of surgery, and were administered desipramine and pargyline (Sigma-Aldrich; 25 mg/kg and 5 mg/kg respectively) intraperitoneally 1 h prior to surgery to increase the selectivity and efficacy of 6-hydroxydopamine (6-OHDA). Upon exposing the previously labeled SNc coordinates on the skull, a small craniotomy was performed and 3 μg 6-OHDA (freshly dissolved in 200 nl of sterile saline containing 0.2% ascorbic acid) was slowly infused at 4.4 – 4.0 mm below dura at a rate of 100 nl/min using a pulled glass micro-pipette. Following surgery, mice were allowed to recover in their cage for at minimum 1 week and were provided with twice-daily intraperitoneally injection of a glucose solution (5% w/v in saline; 1 ml each) and subcutaneous injection of saline (1 ml), and once-daily subcutaneous injections of ketoprofen (10 mg/kg in saline). Water restriction only resumed once mice fully recovered from the lesion and maintained a stable weight.

### Levodopa treatment

Levodopa (Tocris; 1.5-3 mg/kg for Fig. 2d-f, 5 mg/kg for Fig. 2h-j) was administered intraperitoneally along with the peripheral DA decarboxylase inhibitor benserazide hydrochloride (Sigma-Aldrich; 12 mg/kg) in saline. Mice were assayed within 10 min of treatment.

### Fiber photometry

To image rDA1m, we used a custom-made photometry system consisting of a fluorescence mini-cube (Doric; FMC5_E1(460-490)_F1(500-540)_E2(555-570)_F2(580-680)_S) connected to: a) a 565 nm fiber-coupled LED (Thorlabs; M565F3) via a 400 μm, 0.48 NA fiber optic patch cord (Doric), b) a photoreceiver (Newport; 2151) via a 600 μm, 0.48 NA fiber optic patch cord (Doric) and, c) our specimen via a 400 μm, 0.48 NA fiber optic patch cord (Doric). Excitation light was delivered in continuous wave (CW) mode to measure ~30 μW at the patch cord tip. Voltage signals from the photoreceiver were digitized with a National Instruments data acquisition board (PCIe-6353) and breakout terminal block (BNC-2090A) at 2 kHz and recorded with Wavesurfer. To characterize the effects of saline and levodopa treatments on global striatal DA levels, we performed two 30 min-long imaging sessions with the same illumination power and patch cord kept in place: one prior to treatment, and one 10 min after intraperitoneal administration of saline or levodopa. The LED was turned off during the 10 min injection period to avoid photobleaching.

### Optogenetics

To stimulate mDANs, *Dat-ires-Cre::Ai32* mice were chronically implanted with a fiber optic cannula in the right SNc and were connected to a 470 nm fiber-coupled LED (Thorlabs; M470F3) using a patch cord (Thorlabs; M98L01). Light power was adjusted to measure 4 mW at the tip of the patch cord using a power meter (Thorlabs; PM100D and S120C sensor) prior to each experimental session. Optogenetic stimulation (10 ms pulses at 30 Hz for 1 s) was delivered on 30 % of trials (pseudo-randomly arranged) using Wavesurfer starting on trial 11. To control for the effects of light delivery to the brain, sham stimulation sessions were run on separate days using a 595 nm fiber-coupled LED (Thorlabs; M595F2) and the same stimulation parameters, as ChR2 is not excited at this wavelength. To test whether unilateral ChR2 stimulation is sufficiently strong to promote behavioral reinforcement, we placed the same mice for 30 min in a custom-built plastic enclosure fitted on one wall with two nose ports equipped with infrared LED sensors (Digi-key Electronics; 365-1769-ND) to detect head entries. Mice were connected to a 470 nm fiber-coupled LED (Thorlabs; M470F3) via a rotary joint (Thorlabs; RJ1) and patch cord (Thorlabs; M98L01) and ChR2 was stimulated using the same parameters (4 mW, 10 ms pulses at 30 Hz for 1 s) immediately upon head entry into one of the two ports (referred to as the ‘active’ port; counter-balanced across mice) using Wavesurfer. Port entries were recorded using the same program.

### Data processing and analyses

#### Lever push analyses

To determine the speed and amplitude of self-paced lever pushes, we first defined periods of time during which mice pushed the lever vs. not (i.e. ‘active movement epochs’), and then identified individual ballistic pushes in the rewarded direction within these epochs. For the former, we used a method similar to that described by Peters et al. 2014 (ref. ^42^). Briefly, we low-pass filtered the voltage signal provided by the lever’s rotary encoder (down-sampled to 1 kHz) at 10 Hz (4-pole Butterworth), computed the difference between consecutive data points to extract velocity, which we smoothed using a 5-ms moving average window, and extracted the velocity’s envelope using a Hilbert transform. We defined active movement epochs as any time the envelope exceeded a low threshold of 1 mm per second. In well-trained mice, these epochs typically consist of a single, ballistic lever deflection in the rewarded direction. However, they occasionally comprised multiple sequential lever deflections, or instances when mice accidentally let go of the lever mid-push, leading to rapid lever oscillations back to its resting position. To exclude the latter and quantify the speed and amplitude of each goal-directed ballistic push, we isolated individual movement segments within each epoch. To do so, we first obtained higher resolution instantaneous velocity measures using voltage signals provided by the rotary encoder (down-sampled to 1 kHz) low-pass filtered to 30 Hz (6-pole Butterworth) and identified local speed minima. We defined individual movement segments as contiguous periods of time during which speed exceeded 5 mm per second in the rewarded direction. For each movement segment, we calculated the distance traveled by the lever (i.e. amplitude) and the largest instantaneous velocity recorded. To avoid including rare high-velocity artefacts caused by mice letting go of the lever, we restricted analyses to movement segments that precede moments when the lever suddenly travels back to through its resting position and overshoots it by 2.5 mm or more. Figs. 1 and 2 include all qualifying movement segments. Fig 3 reports the maximum deflection achieved (irrespective of the number of movement segments performed) and the maximum velocity recorded per control (i.e. without light stimulation) or stimulation trial. In all cases, session medians for peak velocity and amplitude for each mouse were calculated by bootstrapping (10,000 times). The effects of optogenetic manipulations were calculated as a percent change in peak velocity and maximum amplitude from control ([light–control]*100/control). To quantify changes in lever push speed and amplitude caused by the increase in ‘rewarded push’ threshold, we compared performance for 30 consecutive trials before the threshold change and 30 consecutive trials 200 s after.

#### Photometry

Voltage signals from the photoreceiver were down-sampled to 1 kHz and low-pass filtered at 30 Hz (10-pole, Butterworth). Phasic changes in fluorescence from baseline were quantified as %ΔF/F = (F–F_0_)*100/F_0_ at each time point. We calculated baseline fluorescence (F_0_) by linearly interpolating the lowest 10th percentile of the voltage over 60 s-long sliding windows in 10 s-long steps. For optogenetic experiments, we removed phasic stimulation light artifacts (±6 ms around each light pulse) and filled data by extrapolation using the *inpaint_nans* function (MATLAB Central) prior to filtering. To quantify the power spectrum density (PSD) of DA photometry data, we used the open-source MATLAB package Chronux (http://chronux.org) on unfiltered voltage signal from the photoreceiver and compared power in the 0.5–4 Hz range across conditions (see Fig. 2j) as it is in this range that intrinsic DA fluctuations are most prominent^43^. To estimate changes in global DA levels following intraperitoneal administration of saline or levodopa, we needed to correct for bleaching over time, as well as for any changes in global DA levels that took place during the 10 min that separated the first (pre-injection baseline) and second (post-injection) 30 min-long photometry recordings of rDA1m. To do this, we first plotted baseline F_0_ values during the first imaging session. From this plot of F_0_, we applied a linear fit to the last 10 min to estimate the rate of bleaching (*r_b_*), and calculated the median of the last minute of recording (*F_0_ end value*). The predicted baseline F_0_ values of the 2^nd^ session were calculated as *F_0_ predicted(t)* = *F_0_ end value* + *r_b_* * *F(t)*, starting at *t* = 0 min. We validated this approach with saline-injected mice (Fig. 2h,i). To estimate the peak increase of global DA levels evoked by levodopa, we found the peak of the bleaching-corrected fluorescence in the 2^nd^ session (*F_peak_*) and calculated the percent change as (*F_peak_* – *F_0_ end value*) * *100 / F_0_ end value*.

### Immunohistochemistry

Mice were deeply anesthetized with isoflurane and perfused transcardially with 4 % paraformaldehyde (Electron Microscopy Sciences) in 0.1 M sodium phosphate buffer. Brains were post-fixed overnight and sectioned coronally (50–100 μm) using a vibratome (Leica; VT1000S). Brain sections were mounted on superfrost slides and coversliped with ProLong antifade reagent with DAPI (Molecular Probes). Whole sections were imaged with an Olympus VS120 slide scanning microscope. Tyrosine Hydroxylase (TH) and Dopamine Transporter (DAT) were stained using standard staining protocol (primary antibodies: Mouse monoclonal anti-TH [Immunostar Cat#: 22941 RRID:AB_572268], Rat monoclonal anti-DAT [Millipore Cat#: MAB369 RRID: AB_2190413]; secondary antibodies: Goat anti-mouse IgG Alexa Fluor 647 and Goat anti-rat IgG Alexa Fluor 647 [Thermo Fisher Scientific Cat#: A21236 RRID: AB_2535805 and Cat#: A21247 RRID: AB_141778]). To quantify fluorescence intensity in Fig. 2c, three coronal sections were chosen for each mouse at AP levels +1.5, +1.0 and +0.5 mm from bregma. Regions of interest (ROIs) of the same size were drawn in the left cortex and the left (intact) and right (lesioned) dorsal striatum. The mean fluorescence intensity of DAT in striatum was measured and normalized to that of cortex for comparison.

### Statistics

Normally-distributed data (assessed using Shapiro–Wilk test) were compared using the following parametric statistical tests (as indicated in the text): Student’s paired t-test for comparisons between paired data points, Student’s two-sample t-test for comparisons between un-paired data points, and one-way and two-way analysis of variance (ANOVA) followed by Dunn–Šidák multiple comparison test for comparisons between multiple groups. The following non-parametric tests were used for non-normally-distributed data: Wilcoxon signed-rank test for comparisons between paired data points, Mann–Whitney U test for comparisons between unpaired data points, Kruskal–Wallis one-way analysis of variance, and Friedman test for comparisons across multiple groups. Unbalanced two-way ANOVA were used for comparing animals with multiple experimental conditions with unbalanced repeats (Extended Data Fig. 1e,f). N represents number of mice, unless indicated otherwise. Data points for each mouse in figures reflect bootstrapped session medians. Paired data are linked by gray lines. Summary data across groups are reported in text and figures as mean ± standard error of the mean (SEM), with shaded areas and error bars in figures representing SEM, unless otherwise noted. Exact p-values are provided in text and figure legends, and statistical significance in figures is presented as *, **, *** for p < 0.05, <0.01, <0.001 respectively.

## Acknowledgments

We thank members of the Tritsch laboratory for comments on the manuscript. We thank the Sippy and Schneider labs at NYU for help designing the behavioral apparatus. We acknowledge the New York University Langone Health Genotyping Core Laboratory for mouse genotyping, the Department of Comparative Medicine for animal care and maintenance and the Neuroscience Institute’s imaging facilities for microscope availability. This work was supported by the National Institutes of Health (DP2NS105553 and R01MH130658 to NXT) and the Dana, Whitehall and Parkinson’s Foundations (to NXT).

## Author contributions

HL and NXT conceived of the project, designed and performed experiments, analyzed and interpreted the data, and wrote the manuscript with inputs from all other authors. RM and AS helped perform experiments, RZ assisted with histology, and MM and JRM performed DA sensor imaging in DA-intact and DA-depleted mice.

## Supplementary Information

All data, code, and materials will be made available upon publication. Correspondence and requests for materials should be addressed to NXT.

**Extended Data Fig. 1:**
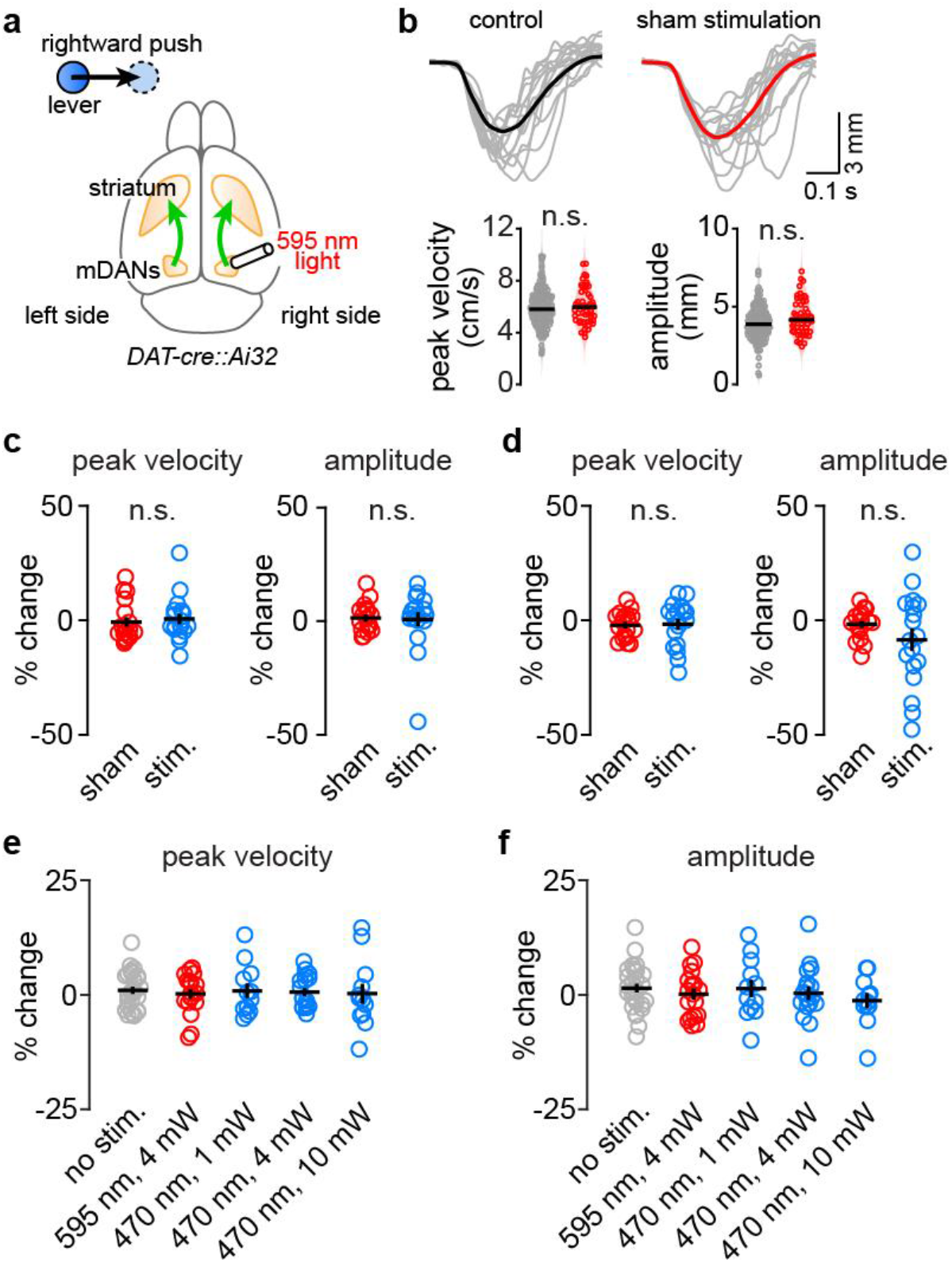
Controlling for different stimulation parameters. **a**, Experimental setup. **b**, *Top*, example lever pushes (session average with 15 individual traces in gray) on trials with (*right*) and without (*left*) 595 nm light (4 mW, 10 ms pulses at 30 Hz for 1 s) in the ventral midbrain. *Bottom*, peak push velocities (p=0.79; Mann–Whitney) and amplitudes (p=0.72, Mann–Whitney) from one session. Black bar: median. **c**, % change in peak push velocity (*left*) and amplitude (*right*) for trials in which 595 nm (sham; red) or 470 nm (ChR2 stimulation; blue) light was delivered in the midbrain compared to interleaved control trials without any light. Analyses were restricted to lever pushes initiated *after* light offset (velocity: p=0.15; amplitude: p=0.86, Friedman; N=18 sessions in 6 mice). **d**, Same as **c** for lever pushes initiated *during* the 1 s-long light stimulation (velocity: p=0.37; amplitude: p=0.48). **e**, % change in peak push velocity on trials with different optogenetic manipulations (10 ms pulses at 30Hz for 1 s at indicated wavelengths and powers) relative to within-session control trials without light. Gray, no light stimulation (randomly-selected 30% of trials in control session). Each circle represents the median value for a session. p=0.98, unbalanced 2-way ANOVA. **f**, Same as **e** for amplitude. p=0.67.

